# A theoretical analysis of single molecule protein sequencing via weak binding spectra

**DOI:** 10.1101/352310

**Authors:** Samuel Rodriques, Adam Marblestone, Ed Boyden

## Abstract

We propose and theoretically study an approach to massively parallel single molecule peptide sequencing, based on single molecule measurement of the kinetics of probe binding [1] to the N-termini of immobilized peptides. Unlike previous proposals, this method is robust to both weak and non-specific probe-target affinities, which we demonstrate by applying the method to a range of randomized affinity matrices consisting of relatively low-quality binders. This suggests a novel principle for proteomic measurement whereby highly non-optimized sets of low-affinity binders could be applicable for protein sequencing, thus shifting the burden of amino acid identification from biomolecular design to readout. Measurement of probe occupancy times, or of time-averaged fluorescence, should allow high-accuracy determination of N-terminal amino acid identity for realistic probe sets. The time-averaged fluorescence method scales well to extremely weak-binding probes. We argue that this method could lead to an approach with single amino acid resolution and the ability to distinguish many canonical and modified amino acids, even using highly non-optimized probe sets. This readout method should expand the design space for single molecule peptide sequencing by removing constraints on the properties of the fluorescent binding probes.

**Author summary:** We simplify the problem of single molecule protein sequencing by proposing and analyzing an approach that makes use of low-affinity, low-specificity binding reagents. This decouples the problem of protein sequencing from the problem of generating a high-quality library of binding reagents against each of the amino acids.

## Introduction

Massively parallel DNA sequencing has revolutionized the biological sciences [2,3], but no comparable technology exists for massively parallel sequencing of proteins. The most widely used DNA sequencing methods rely critically on the ability to locally amplify (i.e., copy) single DNA molecules - whether on a surface [4], attached to a bead [5], or anchored inside a hydrogel matrix [6] - to create a localized population of copies of the parent single DNA molecule. The copies can be probed in unison to achieve a strong, yet localized, fluorescent signal for readout via simple optics and standard cameras. For protein sequencing, on the other hand, there is no protein ‘copy machine’ analogous to a DNA polymerase, which could perform such localized signal amplification. Thus, protein sequencing remains truly a single molecule problem. While true single molecule DNA sequencing approaches exist [7–9], these often also rely on polymerase-based DNA copying, although direct reading of single nucleic acid molecules is beginning to become possible with nanopore approaches [10] that may be extensible to protein readout [11–13]. Thus, the development of a massively parallel protein sequencing technology may benefit from novel concepts for the readout of sequence information from single molecules.

Previously proposed approaches to massively parallel single molecule protein sequencing [14–16] utilize designs that rely on covalent chemical modification of specific amino acids along the chain. Such chain-internal tagging reactions are currently available only for a small subset of the 20 amino acids, and they have finite efficiency. Thus, such approaches would likely not be able to read the identity of every amino acid along the chain.

An alternative approach to protein sequencing [1,17–19] is to use successive rounds of probing with N-terminal-specific amino-acid binders (NAABs) [1]. Recent studies have proposed that proteins derived from N-terminal-specific enzymes such as aminopeptidases [20], or from antibodies against the PITC-modified N-termini arising during Edman degradation [21], could be used as NAABs for protein sequencing. Yet designing or evolving highly specific, strong N-terminal binders to all 20 amino acids (and more if post-translational modifications, e.g., phosphorylation, are considered) is a challenge. Rather than attempting to improve the properties of the NAABs themselves, we will introduce a strategy - which we term “spectral sequencing” - to work around the limitations of existing NAABs and enable single molecule protein sequencing without the need to develop novel binding reagents.

Spectral sequencing measures the affinities of many low-affinity, relatively non-specific NAABs, collectively determining a “spectrum” or “profile” of affinity across binders, for each of the N-terminal amino acids. This profile is sufficient to determine the identity of the N-terminal amino acid. Thus, rather than requiring individual binders to be specific in and of themselves, we will infer a specific profile by *combining measurements of many non-specific interactions*. The spectral sequencing approach measures the single molecule binding kinetics in a massively parallel fashion, using a generalization of Points Accumulation for Imaging in Nanoscale Topography (PAINT) techniques [22,23] to N-terminal amino acid binders.

In what follows, we first derive the capabilities of single-molecule fluorescence based measurement of probe binding kinetics as a function of probe properties and noise sources. We then apply this analysis to the problem of sequencing proteins by measuring profiles of NAAB binding kinetics. Using a range of randomized NAAB affinity matrices as well as an affinity matrix derived directly from the existing measured NAAB kinetics [1], we simulate sequencing of single peptides and obtain 97.5% percent accuracy in amino acid identification over a total observation period of 35 minutes, even in the presence of up to 5% percent error in the instrument calibration and 25% variation in the true underlying kinetics of the binders, due for example to the effects of nonterminal amino acids.

**Fig. 1.**
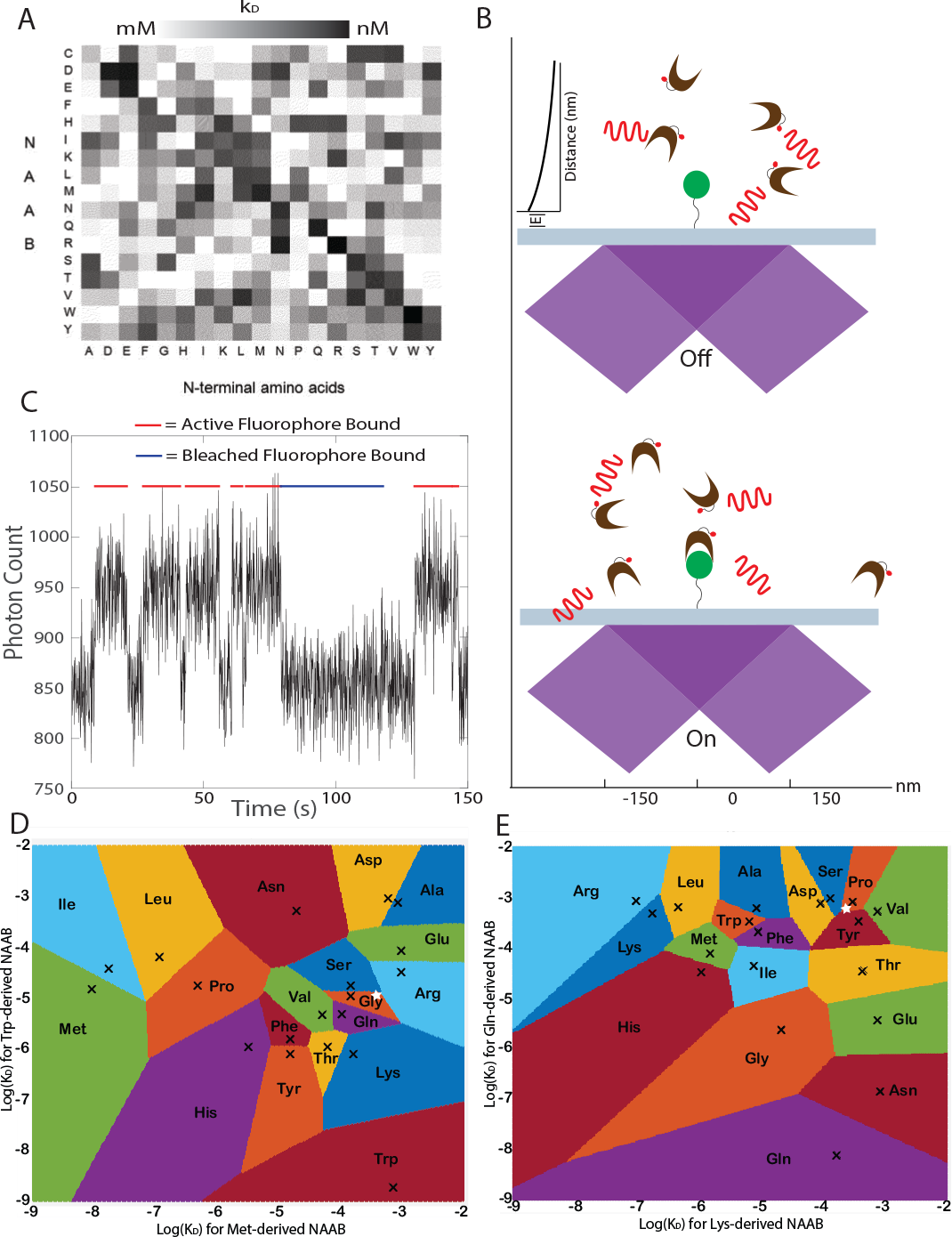
Identifying Amino Acids from Kinetic Measurements. **A** Example affinity matrix for a set of NAABs. The affinities of each of the 17 NAABs are shown for all 19 amino acids excluding cysteine, which is used to anchor the peptides to the surface. Reproduced from [1]. **B** In the proposed measurement scheme, the target (green disk) is attached to a glass slide and is observed using TIRF microscopy. NAAB binders (brown clefts) bearing fluorophores (red dots) are excited by a TIRF beam (purple) and generate fluorescent photon emissions (red waves). **C** When a fluorophore is bound, there is an increase in fluorescence in the spot containing the target. Photobleaching of the fluorophore is indistinguishable from unbinding events, so it is important to use a dye that is robust against photobleaching. Plot shows an illustrative stochastic kinetics simulation incorporating Poisson shot noise of photon emission. **D** The plot shows the affinities of the methionine targeting and tryptophan targeting NAABs for each of the natural amino acids excluding cysteine (black Xs). Upon measuring the affinities for these NAABs against an unknown target, the target can be identified with the amino acid corresponding to the colored region within which the plotted affinities fall. As an example, a pair of measurements yielding the white star would identify the target as glycine. **E** The affinities of the glutamine and lysine targeting NAABs are shown for each of the amino acids. Some amino acids that are practically indistinguishable using the Met and Trp NAABs are easily distinguished using the Gln and Lys NAABs. As an example, if the same target amino acid described in D were measured with only the Gln and Lys NAABs, yielding the white star, we would identify the target as proline. However, combining these measurements with those for the white star in D with Met and Trp NAABs, we see that the true identity of the target is serine. Thus, the higher dimensional measurement of the amino acid using many different NAABs allows disambiguation of the amino acid identity.

## Problem Overview

We consider the problem in which a set of peptides is immobilized on a surface and imaged using total internal reflection fluorescence (TIRF) microscopy. The surface must be appropriately passivated to minimize nonspecific binding [19,24–30]. The limited vertical extent of the evanescent excitation field of the TIRF microscope allows differential sensitivity to fluorescent molecules which are near the microscope slide surface, which allows us to detect NAABs that have bound to peptides on the surface. Existing sets of NAABS (e.g. [1]), derived from aminopeptidases or tRNA synthetases with affinities biased towards specific amino acids, have low affinity or specificity (figure 1A), so one cannot deduce the identity of an N-terminal amino acid from the binding of a single NAAB. Instead, we propose to deduce the identity of the N terminal amino acid of a particular peptide by measuring optically the kinetics of a set of NAABs against the peptide. After observing the binding of each NAAB against the peptide, we will carry out a cycle of Edman degradation [31,32], revealing the next amino acid along the chain as the new N-terminus, and then repeat the process. The process of observing binding kinetics with TIRF microscopy (figure 1B,C) is similar to that used in Points Accumulation for Imaging of Nanoscale Topography (PAINT [22]), e.g., DNA PAINT [23]. This process produces a high-dimensional vector of kinetically-measured affinities at each cycle (figure 1D,E) that can be used to infer the N-terminal amino acid.

This method, while powerful and potentially applicable for current NAABs, ultimately breaks down for probes whose binding is extremely weak, i.e., for which the bound time is so short that only a small number of photons is released while the probe is bound. While fast camera frame rates can be used, the system ultimately becomes limited in the achievable fluorescent signal to noise ratio, unless the measurements are averaged over long experiment times. To extend these concepts into the ultra-weak binding regime, therefore, we propose not to measure the precise binding and unbinding kinetics but rather the time-averaged luminosity of each spot, which indicates the fraction of time a probe was bound. We find that this luminosity-based measurement scheme is highly robust and compatible with short run times.

## Results

Our results are divided into three sections. We first consider the regimes of binder concentration and illumination intensity within which one would expect the proposed method to operate. We then discuss two possible methods for analyzing single molecule kinetic data. Finally, we perform simulations using the derived parameters and data analysis methods in order to estimate the sensitivity of the proposed sequencing method.

### Distinguishability of Amino Acids Based on their NAAB Binding Profiles

A set of binders (NAABs) is characterized by their affinities for their targets (e.g., the 20 amino acids), which can be expressed in the form of an affinity matrix. The affinity matrix *A* is defined such that the *i,j*th entry of *A* is the negative log affinity of the *i*th binder for the *j*th target:

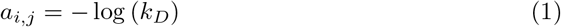

where *k_D_* is the dissociation constant (we define *τ_D_* as the dissociation time).

Throughout this paper, the values of the affinities encoded in the affinity matrix will be referred to as the *reference* values, to distinguish them from the *measured* values obtained in the experiment and from the *true* values, which may depend on environmental conditions but which are not known by the experimenter; the reference values are known and will be used in our computational process of identifying amino acids. As shown in **S1.1 Appendix**, we estimate that it would be possible to determine the identities of the N terminal amino acids from affinity measurements with 99% accuracy, provided that the affinity measurements occur according to a distribution centered on the reference value with standard deviation no greater than 64% of the mean.

### Constraints on Realistic Binding Measurements

In this section, we discuss the primary constraints that are imposed by the measurement modality.

#### Binder Shot Noise

For the purposes of our analysis, we will assume that all binders within 100 nm of the surface emit photons at an equal rate, while more distant binders emit no photons at all. We will also assume that all emitted photons are collected. In reality, excitation due to higher-order beams that do not reflect at the interface will lead to some diffuse background from the bulk solution, and not all photons will be collected due to finite efficiencies in the optical path and at the detector, but contributions from these factors will depend significantly on the specifics of the optical setup and are difficult to estimate; we account approximately for some of these factors in the simulations below by calibrating with published DNA PAINT experiments. We will use the term “observation field” to refer to the region occupied by fluorescent NAABs binding to a single, well-isolated, surface-anchored peptide. For the sake of simplicity, we will assume that the observation field is imaged onto a single pixel on the camera, and will assume that it constitutes a cylindrical region 300 nm in diameter and 100 nm in depth, corresponding to visible TIRF illumination.

In order to be able to distinguish the bound state from the unbound state, the number of photons emitted over the period of observation in the bound state must be significantly larger than the number of photons emitted in the unbound state. We denote by *τ*_obs_ the observation period (which may extend over multiple camera frames), by *R* the rate at which fluorophores in the observation field emit photons, and by *n*_free_ the number of free binders in the observation field, which we will refer to as the “occupation number” for brevity. The occupation number may be given in terms of the volume *V* of the observation field and the molecular number density of the binders *ρ* by

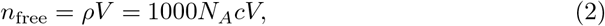

where *c* is the molar concentration and *N_A_* is Avogadro’s number. Then there are two regimes in which we are interested, corresponding to *n*_free_ ≫ 1 and *n*_free_ ≤ 1. The choice of *n*_free_ is up to the experimenter and may be chosen differently for different NAABs. It will need to be optimized to maximize the dynamic range of the *k_D_* readout experiment.

If *n*_free_ ≫ 1, the number of photons emitted by the *n*_free_ free fluorophores in the observation field during the observation period will be drawn from a Poisson distribution with mean and variance

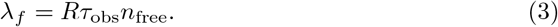

On the other hand, in the bound state, the mean number of photons emitted is

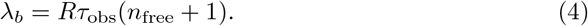

One may then derive (**S1.2 Appendix**) the requirement that

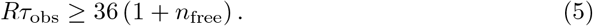

The photon rate *R* is associated with the illumination intensity by

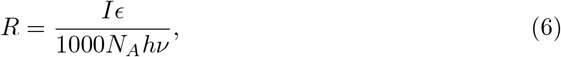

where *ϵ* is the molar absorptivity. (See **S1.3 Appendix** for a derivation.) The minimum intensity that can be used is thus set by the constraints on *R* in equation (5). We obtain

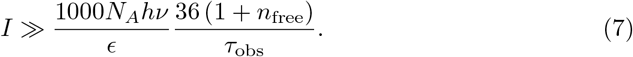

It is worth bearing in mind that an occupation number of *n*_free_ ≈ 1 in every cylinder with diameter 300 nm and height 100 nm corresponds to a molar density of 235 nM.

In the case of *n*_free_ ≤ 1, the noise may deviate significantly from a Poisson distribution (see **S1.2 Appendix** for a discussion). In this regime, it is likely easy to distinguish the bound and unbound states, and instead the constraints on *R* and *τ*_obs_ are set by the requirement that *Rτ*_obs_ be greater than the read and dark noises of the camera. Modern sCMOS cameras have very low dark noises of 0.1 *e*^−^ per second, and read noises of only 1 to 2 *e*^−^ on average. We denote by *p* the per-frame noise, measured in electrons, and by *f* the camera frame rate. Note that *τ*_obs_ may be determined independently of *f*, because the photon counts from multiple frames may be averaged in order to extend the observation period. Instead, *f* is constrained by practical considerations such as the per-frame read noise and the saturation point of the sensor. In order to overcome the read and dark noises, we need

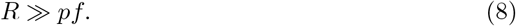

The minimum intensity can thus be determined by the constraint

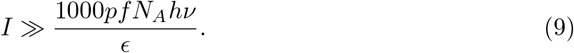

A detector noise of *p* = 1 electron per frame is now standard. To satisfy the requirement in equation (8) for our further calculations, we will take as a requirement that in the limit of *n*_free_ ≤ 1, we should have

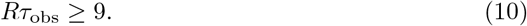

#### Photobleaching

The upper bound on the tolerable intensity is placed by photobleaching. Assuming continuous imaging, the fluorophore should remain active for the entire duration during which the fluorophore is bound. We denote by *N_q_* the average number of photons that a fluorophore emits before it bleaches. Then, we must have

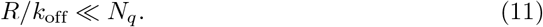

In terms of the intensity,

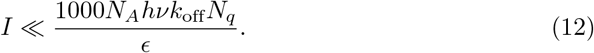

For a typical dye, such as ATTO647N, values of *N_q_* on the order of 10^7^ and *ϵ ~* 1.5 × 10^7^ M^−1^ m^−1^ have been reported [23].

#### Stochastic Binding

Due to the stochastic nature of binding events, the length of the experiment must be chosen to be much longer than the average time between binding events. Hence,

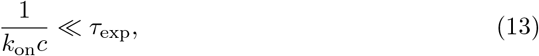

where *c* is the concentration of free binders in the solution.

**Fig. 2.**
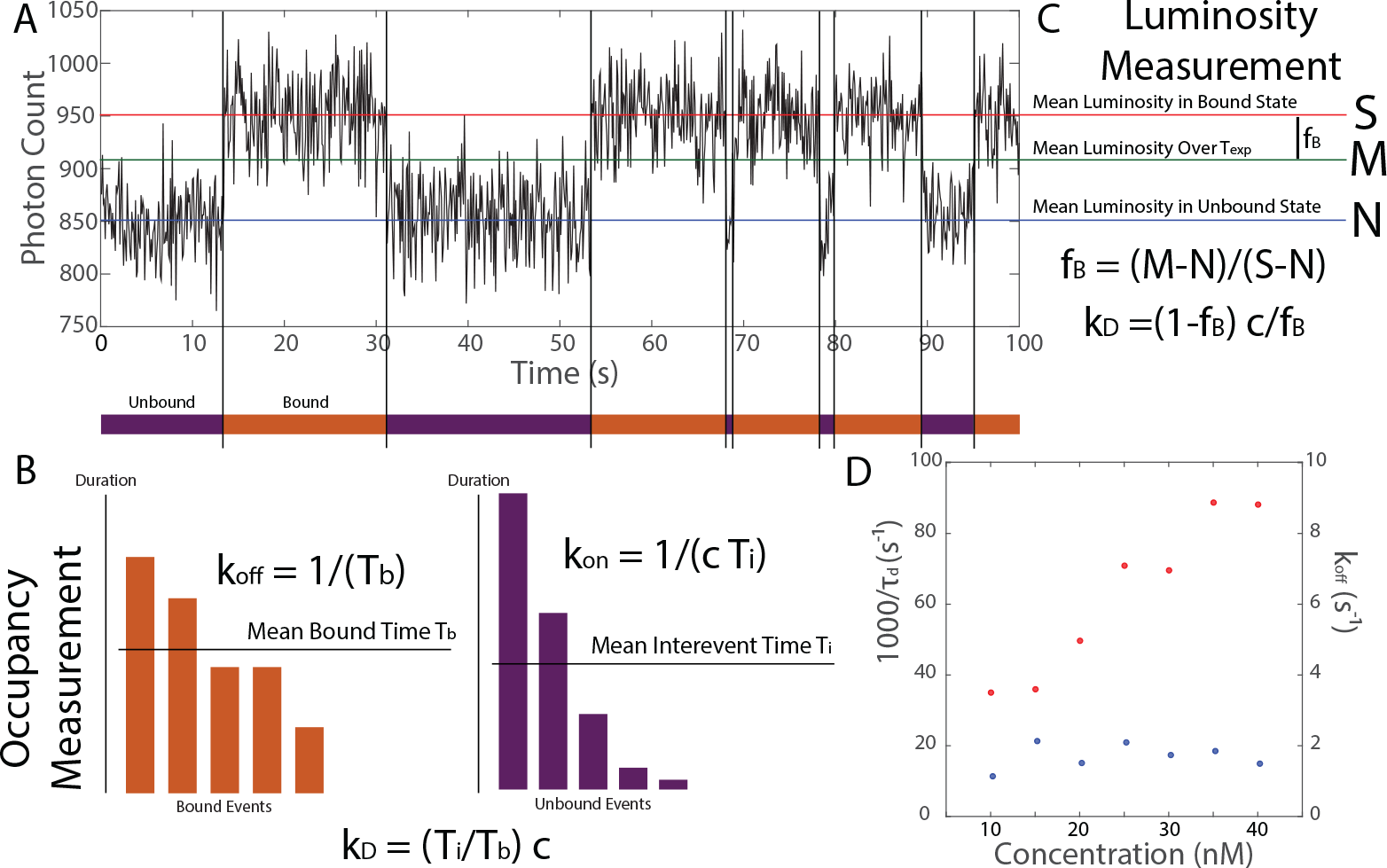
Two Types of Affinity Measurements using TIRF Microscopy. **A** A measurement performed using the proposed scheme yields a fluorescence intensity trace where periods of high intensity correspond to the target being bound and periods of low intensity correspond to the target being free. The affinity of a binder against the target may then be determined in two ways, either via occupancy measurements or via luminosity measurements. **B** An occupancy measurement is performed “along the time axis,” by calculating *k*_on_ from the average time between binding events, and *k*_off_ from the average length of binding events. **C** On the other hand, a luminosity measurement is performed “along the brightness axis,” by calculating *k_D_* directly from the average luminosity of the target over the whole observation period. **D** We validated our simulation by applying occupancy measurements to determine kon and koff from simulated data. The parameters used here were identical to those used in the production of Figure 2a in [23]. See text for symbol definitions.

## Methods of Data Analysis

A measurement performed using this scheme yields a time series such as that shown in Figure 2A. We now discuss the two primary options for extracting the kinetics from this data and the experimental conditions that are optimal for each scheme, given the constraints discussed above.

### Occupancy Measurements

The first measurement, used commonly in the field of single-molecule kinetics [23,33], relies on detecting changes in the occupancy state of the target. The measurement scheme is depicted schematically in Figure 2B. This measurement is performed “along the time axis,” in the sense that it relies on temporal information - *when* probes bind and unbind - and is relatively insensitive to analog luminosity information beyond that needed to make these digital determinations. This method is optimal for measurements on binders with very high affinities, which can be performed at low concentrations. The upper limit on the dynamic range of this method is set by the frame rate, i.e.,

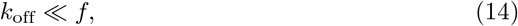

where *f* is the imaging rate. In order to extract temporal information, we set *τ*_obs_ = 1/*f*. This method will typically operate in the limit *n*_free_ ≤ 1, so from equation (10), we find that we must have *Rτ*_obs_ ≥ 9. Hence,

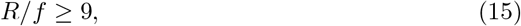

and hence

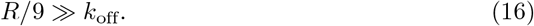

On the other hand, the lower bound on the dynamic range is provided by photobleaching, as captured in equation (11). In total, we have

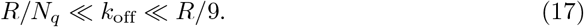

In practice, for this measurement modality, we will choose *f* = 100 Hz and *R* = 10^4^ s^−1^, corresponding to a laser power of 13 Wcm^−2^. With *N_q_* ~ 10^7^, the requirement becomes *k*_off_ ≪ 100 s^−1^ and *k*_off_ ≫ 10^−3^ s^−1^, yielding an effective dynamic range of approximately three orders of magnitude of *k*_off_.

Finally, the experiment time is constrained by the requirement that

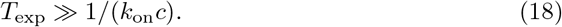

For a value of *k*_on_ on the order of 10^5^ M^−1^ s^−1^ and a concentration on the order of 100 nM, this requirement implies that an experiment time of at least 100 seconds is necessary in order to see several binding events with high probability.

If the binding and unbinding events may be identified, then one may determine the average binding time *T_b_* and the average time between binding events *T_i_*, which we will refer to as the inter-event time. If photobleaching may be neglected, then we have

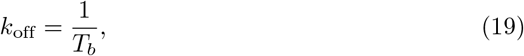

and

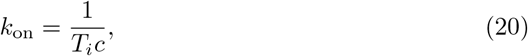

where *c* is the free binder concentration. Thus,

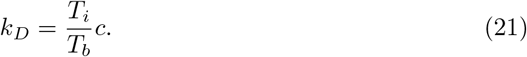

Alternatively, if the on-rate *k*_on_ is known, then it is possible to determine koff even in the presence of photobleaching. (See **S1.4 Appendix** for details.)

### Luminosity measurements

An alternative to the occupancy-time measurements described above involves deducing *k_D_* directly from the *fraction f*_*B*_ of time that the target is bound by a probe. This quantity may in turn be deduced from the *average* luminosity of the spot containing the free binder over the period of observation, as depicted in Figure 2C. Whereas occupancy measurements are performed “along the time axis,” neglecting luminosity information, luminosity measurements are performed “along the luminosity axis,” neglecting temporal information about the series of binding and unbinding events. Because it does not attempt to track individual binding and unbinding events, this method is particularly suited to measurements of weak binders performed at high background concentrations, where binding and unbinding events may occur faster than the camera frame rate. Moreover, this method relies on each NAAB of a given type having approximately the same brightness, which could be achieved using a high-efficiency method for monovalently labeling the NAAB N- or C-terminus [34,35].

If the target is bound a fraction *f_B_* of the time, then the dissociation constant is given by

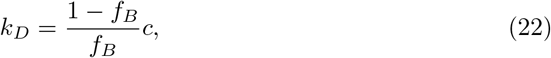

where *c* is the background binder concentration. We denote by *S* the average brightness of the spot when a fluorescent binder is attached to the target, and by *N* the average brightness of the spot when the target is free. Neglecting photobleaching, the average brightness of the spot over the whole experiment is given by

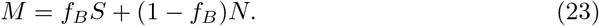

If *S* and *N* are known, then *f_B_* may thus be deduced directly from the measured photon rate *M* averaged over the entire experiment, via

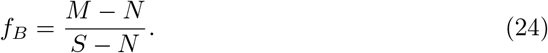

*S* and *N* can be measured directly for example by anchoring NAABs sparsely to a surface and measuring the brightness of the resulting puncta (to deduce *S*), or puncta-free regions (to measure *N*).

One significant advantage of this method is that the observation period *τ*_obs_ can be chosen to be arbitrarily long by averaging the photon counts of many successive frames (i.e., we have *τ*_obs_ = *T*_exp_). In practice, we will use *τ*_obs_ = 100s. With this value, we can use a relatively high concentration of 2 μM (corresponding to *n*_free_ ≫ 1) even for a relatively low intensity of 1.3W cm^−2^ (corresponding to *R* = 10^3^ s^−1^, while still satisfying (5). Operating in this regime significantly reduces the vulnerability of the experiment to stochasticity and photobleaching. However, unlike in the case of occupancy measurements, there is no way to account for photobleaching, if it occurs. Nonetheless, we do not expect photobleaching to have a significant impact on our results, since most of the NAABs have fairly high off-rates [1,20].

In contrast to occupancy measurements, luminosity measurements are also sensitive to error in the calibration of the measurement apparatus, for example if the brightness of the bright and dark states is not known exactly. The bright and dark states *S* and *N* could likely be calibrated by doping in labeled reference peptides to the sample to be sequenced. Still, there may be some error in the measurements of *S* and *N*. For a discussion of computational strategies for coping with calibration error, see Appendix 1.5.

## Simulations

### Simulation Outline

In order to determine whether the TIRF measurement scheme described above can be used to identify single amino acids on the *N*-termini of surface-anchored peptides, we simulated N-terminal amino acid identification experiments.

We first used a specific NAAB affinity matrix given in [1]. Importantly, random affinity matrices generated by permuting the values of the NAAB affinity matrix perform similarly well in residue-calling simulations (fig 5 and 6). To generate the random affinity matrices with statistics matching the statistics of the NAAB affinity matrix, each matrix element was chosen by randomly sampling values from the NAAB affinity matrix of [1], without replacement. The simulations described here can therefore be assumed to apply to general ensembles of N-terminal binders with affinity value statistics similar to those displayed by these existing NAABs.

In the simulations, there is assumed to be one free target in the volume analyzed, which is a cylinder of diameter 300 nm and height 100 nm as discussed above. The simulation considers each frame of the camera in succession, and models the number of photons registered at the camera. At the start of the simulation, or as soon as the target becomes free, a time *T*_free_ is drawn from an exponential distribution with mean 1/(*k*_on_*c*), where *c* is the concentration of binders. Once a time equal to *T*_free_ has passed, the binder is considered occupied, and a time *T*_bound_ is drawn from an exponential distribution with mean 1/*k*_off_. In addition, upon binding, a time *T*_photobleach_ is drawn from an exponential distribution with mean *N_q_/R*, where *N_q_* is the number of photons the fluorophore emits on average before bleaching and *R* is the single-fluorophore photon rate. If the time *T*_photobleach_ is less than the time *T*_bound_, the fluorophore ceases to emit photons after time *T*_photobleach_. Within a given frame, the simulation tracks binding, unbinding, and photobleaching events, and computes the number of signal photons detected by the camera by drawing from a Poisson distribution with mean *RT*_on_, where *R* is the single fluorophore photon rate and *T*_on_ is the amount of time during the frame in which an unbleached fluorophore was attached to the target.

The dominant contribution to noise in the simulation is expected to come from fluorophores attached to free binders that enter and leave the observation field [33]. At the end of each frame, the simulation draws the number of free binders that enter the observation field during the frame from a Poisson distribution with mean *n*_free_/*f*, where *f* is the frame rate and *n*_free_ is the free binder occupation number of the frame. For each binder that enters the observation field, we draw a dwell time *t* from an exponential distribution with mean *τ*_dwell_ as calculated in equation (40) from diffusion theory (see Appendix 1.2), and a total photon contribution from a Poisson distribution with mean *Rt*. Finally, we calculate the detector shot noise from a Gaussian distribution with mean *p* and standard deviation equal to 0.1*p*.

#### Validation of the Simulation Pipeline

To validate the simulations, we reproduced the DNA PAINT kinetics data collected by [23] using the parameters reported in that paper. There, values of *k*_on_ = 10^6^ M^−1^ s^−1^ and *k*_off_ = 2 s^−1^ were reported. Imaging was conducted at 650 nm with a power of 4 mW to 8 mW over an imaging region of (150μm)^2^, corresponding to an intensity of approximately 26.67 Wcm^−2^, corresponding to a photon rate of *R* ~ 18 000 s^−1^, assuming a dye comparable to ATTO655. However, accounting for the low quantum efficiency of ATTO655 and possible losses of light in the light path of the microscope, we performed our simulations with *R* ~ 1500 s^−1^. From our simulated data, we were able to reproduce the measured off- and on-rates, as shown in Figure 2D. Moreover, consistent with [23], photobleaching only became apparent in the simulation at laser powers greater than 100 mW.

### Measurements of *k_D_*

#### Occupancy Measurements

We next simulated occupancy measurements of the binding kinetics of the NAAB against the target. We performed 100 simulations for each of five different values of *k*_on_ between 10^4^ M^−1^ s^−1^ and 10^6^ M^−1^ s^−1^, which is consistent with standard values observed for antibodies [36], and for each of five different values of *k_D_* between 100 μM and 10 nM. We assumed a framerate of 100 Hz, detector read noise of 1 e^−^, and a laser power of 130 kW m^−2^, corresponding to a single-fluorophore photon rate of 10^4^ s^−1^. NAABs were washed onto the sample at a concentration of 300 nM, and each wash was observed for *T*_exp_ = 100 s.

**Fig. 3.**
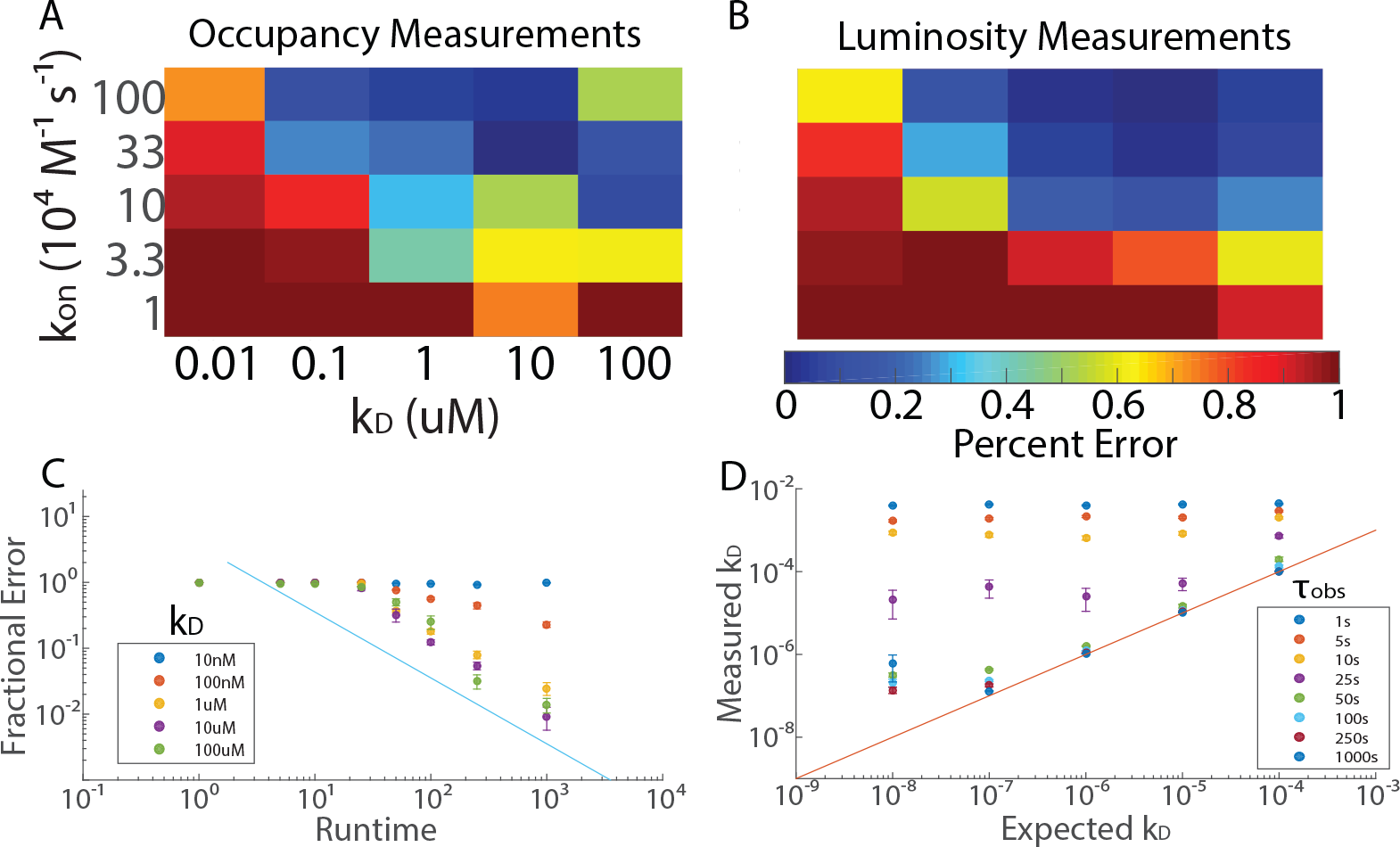
Two Types of Affinity Measurements using TIRF Microscopy. **A** The accuracies of occupation measurements of *k*_*D*_ are shown as a function of *k*_*D*_ and *k*_on_ for the simulation described in the text, with *T*_exp_ = 100s. These measurements achieve high accuracy for *k*_on_ ≥ 10^5^ M^−1^ s^−1^ and *k*_off_ ≪ 100 s^−1^. For values of *k*_off_ on the order of 100s^−1^ (upper right-hand corner), the accuracy deteriorates significantly. **B** The accuracies of luminosity measurements of *k*_*D*_ are shown as a function of *k*_*D*_ and *k*_on_. These measurements achieve high accuracy for *k*_on_ ≥ 10^5^ M^−1^ s^−1^ and *k*_*D*_ ≥ 100 nM. The heat map shown gives the fractional errors as a function of *k*_*D*_ and *k*_on_ for the simulation described in the text, with *T*_exp_ = 100 s. In contrast to occupation measurements, the accuracy of luminosity measurements does not deteriorate for very high values of *k*_off_. **C** For luminosity measurements only, the mean fractional error in the measured value of *k*_*D*_ is plotted as a function of the observation time for five different values of *k*_*D*_. The line *y* = 1/*x* is plotted as a guide to the eye. For *k*_*D*_ = 10 nM and *k*_*D*_ = 100 nM, the effects of photobleaching are evident at longer runtimes. **D** Also, for luminosity measurements only, the measured value of *k*_*D*_ is plotted as a function of the actual value of *k*_*D*_ for 8 different values of the runtime. The performance of the algorithm improves dramatically for *τ*_obs_ > 25 s. The line *y* = *x* is plotted as a guide to the eye. Error bars in C, D denote standard error over 100 trials.

In order to analyze the data, we ran a control simulation in which *k*_on_ was set to 0, so that no NAABs bound to the target. In practice, this calibration could be performed by observing a spot that does not have a target. From this, we calculated the mean and standard deviation of the noise on a per-frame basis. We then identified binding and unbinding events as follows. First, we identified all frames in which the photon count was more than 2 standard deviations above the noise mean. These frames will be referred to as “on” frames, whereas all other frames will be referred to as “off” frames. If three such “on” frames occurred in a row, the event was identified as a binding event. The binding event was considered to continue until at least two “off”-frames in a row were observed. Once all the binding and unbinding events were identified, the average inter-event time and the average binding time were calculated, and from these the kinetics were deduced (Figure 2A).

The accuracy of the *k_D_* measurements was found to improve with increasing *k*_on_, and to improve with increasing *k_D_* for values of *k*_off_ below 10 s^−1^ (Figure 3A). For values of *k*_off_ significantly above 10 s^−1^, it was no longer possible to distinguish individual binding and unbinding events from noise (Figure 3A, upper right-hand corner). Moreover, for values of *k*_on_ below 10^5^ M^−1^ s^−1^, the condition *T*_exp_ ≫ 1/(*k*_on_*c*) was no longer satisfied. Finally, for very small values of *k_D_*, photobleaching limited the accuracy of the analysis. For *k*_on_ > 10^5^ M^−1^ s^−1^ and *k*_off_ ~ 10 s^−1^, it was possible to obtain the correct value of *k_D_* to within approximately 5 − 10%. However, the accuracy deteriorated sharply for combinations of *k*_on_ and *k*_off_ deviating from these ideal conditions.

#### Luminosity Measurements

We then simulated luminosity measurements of *k_D_* using comparable parameters. Because these measurements depend only on the average luminosity over the entire experiment, the entire experiment was lumped into a single camera frame. In practice, however, the same results can be obtained by averaging over the photon counts of multiple frames. The laser intensity was set to 13 kW m^−2^, corresponding to a single-fluorophore photon rate of *R* = 1000 s^−1^, and the free binder concentration was set to 2 μM. The photon rate of the off-state was determined first by running the simulation with the value of *k*_on_ set to 0. The photon rate in the on-state was then determined by running the simulation with the value of *k*_on_ set to 10^10^ M^−1^ s^−1^, and the value of *k_D_* set to 10^−20^ M. Because the exposure time used in this experiment is very long compared to the dwell time of free binders in the observation field, it was assumed that all free binders that enter the observation field emit a number of photons equal to Rτ_dwell_ (i.e., the noise was taken to be approximately Poissonian), which substantially reduces the computational complexity of the algorithm. Once the average luminosity over the experiment was determined, the value of *f_B_* was deduced.

For observation times shorter than 50s, the analysis sometimes returns values of *f_B_* arbitrarily close to or greater than 1 or arbitrarily close to or less than 0. This can happen as a consequence of statistical error in the luminosity measurements, even in the absence of systematic error. For this reason, in order to avoid negative or outlandishly large values of *k_D_* from compromising the analysis, we chose the maximum value of *f_B_* to be equal to the value expected when *k_D_* = 1 nM, and we chose the minimum value of *f_B_* to be equal to the value obtained when *k_D_* = 10 mM. Any values of *f_B_* outside of this range were adjusted to the maximum or minimum value, appropriately.

In order to enable comparison to the occupancy measurements, the simulation was run 100 times for each of five values of *k*_on_ between 10^4^ M^−1^ s^−1^ and 10^6^ M^−1^ s^−1^ and for each of five values of *k_D_* between 100 μM and 10 nM. The accuracy was found to be comparable to that obtained in the occupancy experiments (Figure 3A), except that the accuracy did not deteriorate for very high values of *k*_off_ (Figure 3B, upper right-hand corner). For values of *k*_on_ on the order of (or greater than) 10^5^ M^−1^ s^−1^ and values of *k_D_* greater than 1 μM, *k_D_* could easily be determined to within the accuracy condition required by equation (31).

To ascertain the effect of *τ*_obs_ on the accuracy, the simulation was run 100 times for each of the same 25 combinations of *k*_on_ and *k*_off_, with 8 different values of *τ*_obs_ between 1s and 1000 s and a free binder population of 2 μM (Figure 3C). As expected, the accuracy was found to undergo a sharp transition when *τ*_obs_ was on the order of 25 s, corresponding to 1/(*k*_on_*c*) ≪ *τ*_obs_. For values of *τ*_obs_ > 25 s and values of *k_D_* greater than 1 μM, the error in the measurement of *k_D_* decreased like 1/τ_obs_ (Figure 3C). For observation times greater than 25 s, the value of *k_D_* could be calculated with standard deviation less than 64% of the mean for values of *k_D_* on the order of or greater than 1 μM, although photobleaching leads to saturation and significant losses of accuracy for smaller values of *k_D_* (Figure 3D).

Separately, to ascertain the effect of the free binder concentration on the accuracy, the simulation was run 1000 times on each of the same 25 combinations of *k*_on_ and *k_D_*, with *τ*_obs_ = 50 s at seven different values of the concentration between 10 nM and 5 μM. For values of *k*_on_ such that *τ*_obs_ ≫ 1/(*k*_on_*c*), the effect of increasing *k*_on_ was found to be similar to the effect of increasing *τ*_obs_ (data not shown).

### Identifying Amino Acids

Because standard deviations in *k_D_* below 64% of the mean could consistently be achieved in the luminosity measurements across a broad range of values of *k*_on_ and *k_D_*, it is reasonable to expect that luminosity measurements of NAAB binding kinetics with the affinity matrix in figure 1a could allow for the identification of amino acids at the single molecule level. We thus simulated an experiment in which a peptide with an unknown amino acid is attached to a surface, and is observed successively in multiple baths, each containing a single kind of fluorescent NAAB. In this simulation, amino acids were randomly chosen from a uniform distribution. Binders were added to the solution at a concentration of 1 μM and the laser power was set to 13 kW m^−2^. For each NAAB, effective values of the dissociation constant *k̃_D_*, the on-rate *k̃*_on_, the effective brightness *R̃*, and the calibration levels *S̃* and *Ñ* were determined for the NAAB-amino acid pair. The spot containing the NAAB was then observed over a period of time *τ*_obs_, which ranged from 50 to 500 seconds, and the total number of photons observed was stored. This process was repeated for each NAAB, generating a vector 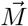 of observed photon counts.

Systematic error in the experiment was parametrized using three quantities. For each NAAB, the effective dissociation constant *k̃_D_* for the NAAB-amino acid pair was drawn from a normal distribution centered on the reference value *k_D_*, with standard deviation equal to *σ_K_ k_D_*, where *σ_K_* parametrizes the effect of non-terminal amino acids and other environmental factors on the dissociation constant. Likewise, the effective brightness of the NAAB relative to the average NAAB brightness was determined by drawing *R̃* from a normal distribution with mean *R* and standard deviation *σ_B_R*, where *R* is the photon rate of a standard fluorophore (assumed here to be ATTO647N) in the observation field. Finally, in order to determine the effective calibration levels, the true calibration levels *S* and *N* were first determined as the luminosity of the bound and unbound states, as described above (Luminosity Measurements). The measured calibration levels *S̃* and *Ñ* were then determined by drawing from a normal distribution with mean equal to *S* and *N* and with standard deviation equal to *σ_C_S* and *σ_C_N*, respectively. The values of *σ_K_*, *σ_B_*, and *σ_C_* will be given below in percentages.

Analysis was performed by comparing the measured photon counts to the photon counts that would have been expected for each amino acid, as described above. For each NAAB-amino acid pair, the expected photon count was calculated from the NAAB concentration *c*, the reference value of *k_D_* and the measured calibration level *S̃* and *Ñ*, via

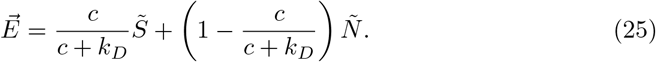

The resulting expected photon counts were then assembled into a matrix *W*, such that the (*i, j*)th element of *W* is the photon count that one would have expected on the measurement of the *i*th NAAB if the target were the *j*th amino acid, given the calibration levels *S̃* and *Ñ*. Finally, the amino acid identity *I_aa_* was determined by minimizing the norm between the vector of observed photon counts 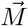 and the columns of *W*, i.e.,

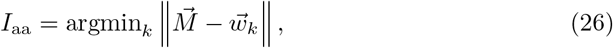

where 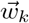 is the *k*th column of *W*.

In Figure 4A-C, the accuracy with which amino acids can be identified is shown as a function of the observation time and the systematic error, for a 1 μM free binder concentration. In the absence of systematic error, amino acids could be identified with greater than 99% accuracy after a 50 s observation. Moreover, if the calibration error can be kept below 5%, and if the systematic error in the kinetics can be kept below 25%, then our simulations indicate that it would be possible to identify amino acids with greater than 97.5% accuracy over an observation window of 100 s.

The measurement accuracy was shown to be robust against systematic differences in brightness between different NAABs (data not shown). The experiment also showed robustness against systematic deviation in *k_D_* up to the 25% level, with progressive deterioration in the measurement accuracy observed for values of *σ_K_* above 25%. Calibration error was found to have the most substantial effect on the accuracy, with calibration errors on the order of 10% reducing the achievable accuracy below 90% even for an observation time of 250 s. The effects of calibration error on the accuracy could be substantially reduced by reducing the concentration of free binders (Figure 4D), which has the effect of increasing the gap between the *S* and *N*. However, in order to preserve the requirement that *T*_exp_ ≫ 1/(*k*_on_*c*), it is necessary to increase the experiment length by a similar factor. (It is worth noting that for this reason, a free NAAB concentration of 1 μM was used, rather than 2 μM as used above.) Moreover, this improvement comes at the cost of increased sensitivity to systematic error in *k_D_*.

## Application to randomized affinity matrices

In order to determine whether the protein sequencing method proposed here is limited to the specific affinity matrix given in [1], we generated affinity matrices with comparable binding statistics by randomly shuffling the *k_D_* values in the NAAB affinity matrix. For 100 such random affinity matrices, we then performed identical simulations as in fig 4E, assuming 5% calibration error and 25% kinetic error. To calculate the overall error rate for a given matrix, we summed the frequencies of incorrect residue calls (the off-diagonal elements of the matrices in fig 4E). The overall error rate for the NAAB affinity matrix, calculated in this way, is 0.0124, and the distribution of error rates across the random matrices is shown in fig 5. Only one randomly generated affinity matrix had an error rate lower than the NAAB error rate. Nonetheless, it is clear that most affinity matrices with affinity statistics similar to the NAABs [1] would yield errors in the range of 1%-4%, and thus the sequencing method described here is generalizable to a range of similar N-terminal amino acid binders.

**Fig. 4.**
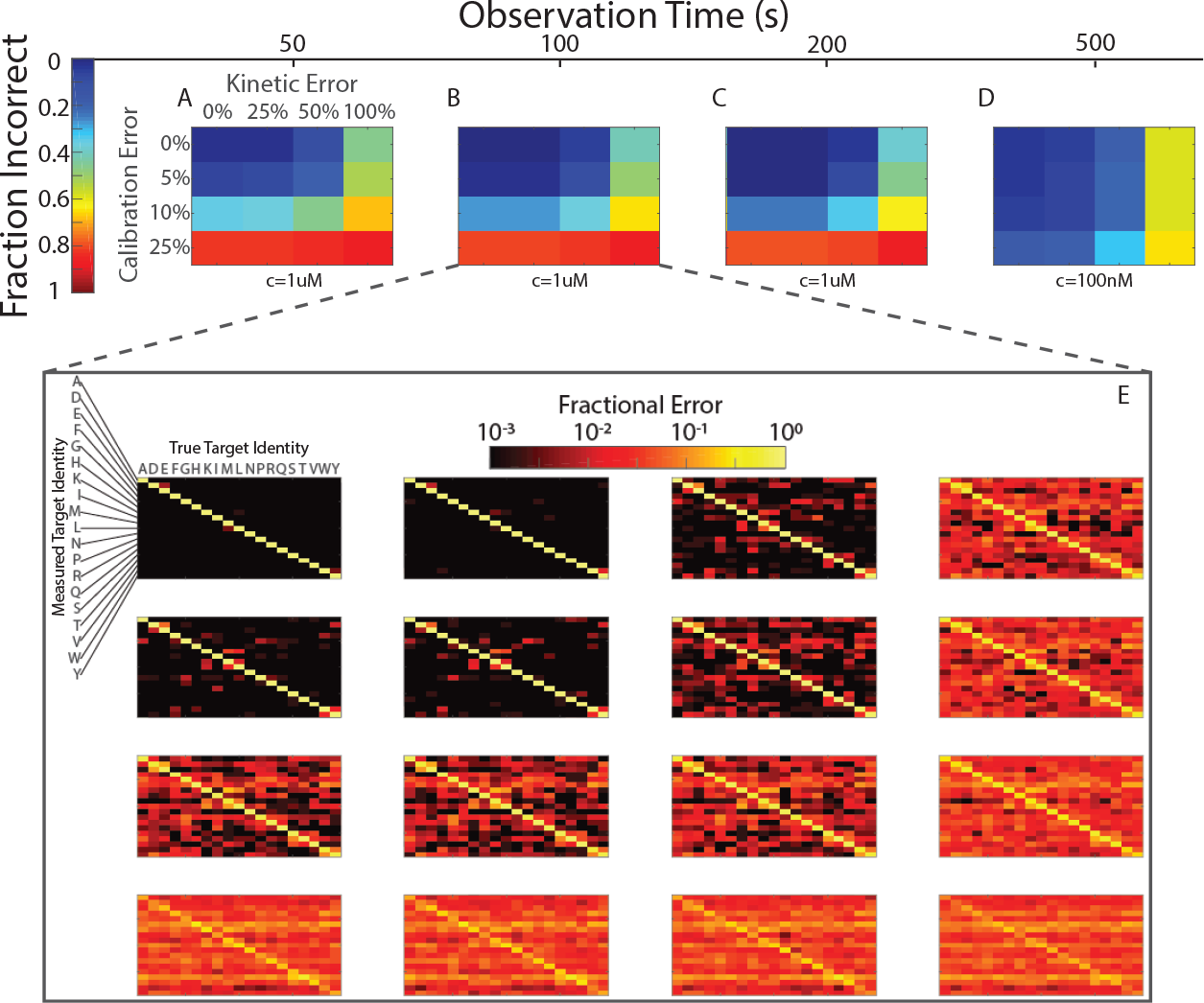
Identification of Amino Acids is Robust Against Systematic Error. The fraction of amino acids incorrectly identified is plotted as a function of *τ*_obs_ for four different values of the systematic calibration error *σ_C_* and four different values of the systematic kinetic error *σ_K_* (as described in the text). **A** In the absence of systematic error, measurements with *τ*_obs_ = 50 s result in correct amino acid identification more than 98% of the time. For 25% error in *k_D_*, the accuracy drops to 97.5%, and if 5% calibration error is added, it drops further to 92%. More than 5% systematic error in the calibration leads to very significant numbers of mistakes in amino acid identification. **B** With *τ*_obs_ = 100 s, an accuracy of 97.5% was obtained for 25% error in *k_D_* and 5% error in the calibration. **C** Increasing *τ*_obs_ beyond 100 s at the same binder concentration leads to diminishing improvements in the accuracy. **D** The sensitivity to calibration error could be substantially reduced by decreasing the concentration of free binders to 100 nM. However, this increased concentration necessitates a longer runtime. **E** For *τ*_obs_ = 100 s, plots are shown for each value of *σ_C_* and *σ_K_*, depicting the probability that a given target amino acid (on the horizontal axis) was assigned a particular identity (on the vertical axis). Off-diagonal elements correspond to errors.

**Fig. 5.**
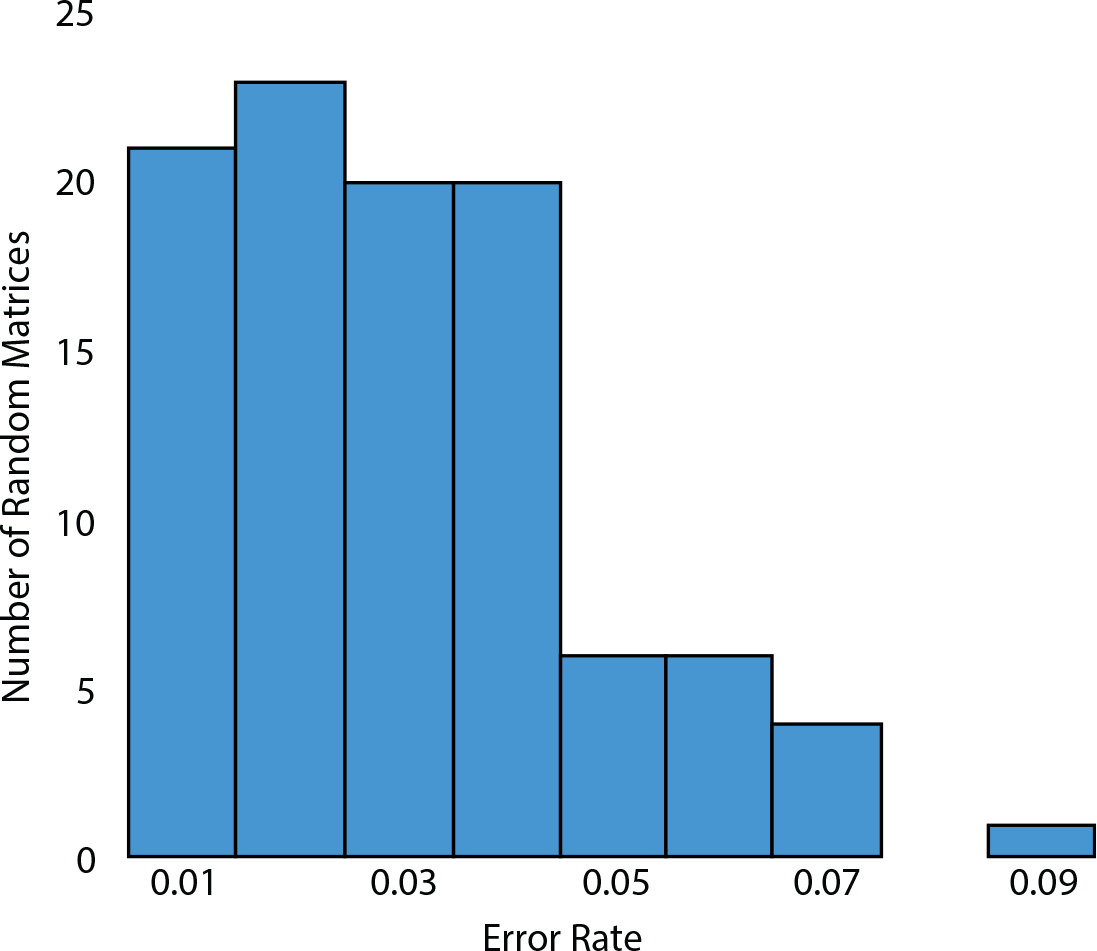
Overall Error Rates for 100 Random Affinity Matrices. The overall error rate, calculated as the sum of incorrect residue calls divided by the total number of residue calls over 10000 trials, is plotted for 100 random affinity matrices.

## Discussion

The calculations and simulations discussed above indicate that if the measurement apparatus can be calibrated with an accuracy of 5%, and if the reference values of *k_D_* can be kept within 25% of the true values, it is theoretically possible to determine the identity of an N-terminal amino acid with greater than 97.5% accuracy by measuring the kinetics of the NAABs against the target amino acid. Crucially, *k_D_* can be inferred just from the time-averaged local concentration of NAABs within the observation field, and thus the measurement can be performed at relatively high background binder concentrations, because it does not rely on being able to distinguish individual binding and unbinding events.

### Primary Uncertainties

Three primary uncertainties exist regarding the validity of the simulations performed here. Firstly, our simulation did not incorporate the effects of non-specific binding of NAABs to the surface. Nonetheless, if such non-specific binding occurs with sufficiently low affinity, we anticipate that the effect of the non-specific binding will be comparable to the effect of increasing the affinity of the binders for the target, and we have shown that our experiment displays considerable robustness against such sources of systematic error. On the other hand, if non-specific binding occurs with high affinity, we anticipate that by examining the time-course of the luminosity, such non-specific binding events can be identified and accounted for.

In addition, some uncertainty exists surrounding the value of *N_q_* for the organic dyes of interest to us, with values between 10^5^ and 10^7^ being reported [23,37]. However, we expect our method to be relatively robust to photobleaching due to the relatively low affinity and high off-rates of most of the NAABs. Moreover, it is possible that more photostable indicators such as quantum dots could be used in place of organic dyes. Note that with any labeling scheme, there will be some concentration of “dark NAABs” that are not labeled. Thus, the concentrations reported for the simulations above should be regarded as the concentrations of “bright NAABs.” The presence of dark NAABs is unlikely to affect the experimental results provided the total NAAB concentration is less than the dissociation constant (i.e., as long as the target is free most of the time), so a high concentration of dark NAABs can always be compensated for by reducing the total NAAB concentration and increasing the measurement duration.

### Parallelization

We anticipate that the approaches discussed here could be parallelized in a way reminiscent of next-generation nucleic acid sequencing technologies, allowing for massively parallel protein sequencing with single-molecule resolution. In the ideal case, if a 64 megapixel camera were used with one target per pixel, we would have the ability to observe the binding kinetics of NAABs against approximately 10^7^ protein fragments simultaneously. With an observation time of 100 seconds per amino acid-NAAB pair, this corresponds to approximately 35 minutes of observation time per amino acid, or 5 days to identify a protein fragment of 200 amino acids in length. On average, therefore, the sequencing method would have a throughput of approximately 20 proteins per second.

However, the throughput of the device could be improved dramatically if the readout mechanism were electrical, rather than optical. CMOS-compatible field-effect transistors have been developed as sensors for biological molecules [38–41]. Moreover, electrical sequencing of DNA has been accomplished using ion semiconductor sequencing [42]. Most recently, CMOS-compatible carbon nanotube FETs have been shown to detect DNA hybridization kinetics with better than 10 ms time resolution [43,44]. Similar CMOS-compatible devices have been adapted to the detection of protein concentrations via immunodetection [45]. These systems have the added benefit that they sense from a much smaller volume than TIRF does (sometimes as small as ~ 10 cubic nanometers [44]), substantially reducing the impact of noise on the measurement. A single 5 inch silicon wafer covered in transistor sensors at a density of 16 transistors per square micron would be capable of sequencing 10^12^ proteins simultaneously, corresponding to an average throughput of 2,000,000 proteins per second on a single wafer, or one mammalian cell every 7 minutes. Such an approach could make use of dedicated integration circuitry to compute the average NAAB occupancy at the hardware level, greatly simplifying data acquisition and processing. Moreover, if the devices were made CMOS-compatible, they could be produced in bulk, greatly improving scalability. If the intrinsic contrast provided by the NAABs is insufficient for measurements with FETs, the NAABs can be further engineered to have greater electrical contrast, for example by conjugating them on the C-terminus to an electrically salient protein such as ferritin. A combination of electrical and optical readouts may also be desirable. Recently, CMOS-compatible single-photon avalanche diode imaging systems have been developed that are capable of detecting the presence of fluorophores on a surface without magnification [46].

Finally, although the use of TIRF microscopy in the case studied here restricts the proposed approach to operate close to a reflecting surface, the use of thin sections or alternative microscopies could potentially allow such protein sequencing methods to operate *in-situ* inside intact cells or tissues.

## Conclusion

We have shown that single molecule protein sequencing is possible using low-affinity, low-specificity binding reagents and single molecule fluorescent detection. Achieving a high-quality single molecule surface chemistry and TIRF measurement setup will be a challenge, but if this can be achieved, our results show that a wide range of binding reagent families should be adaptable to single molecule protein sequencing.

## 1 Supporting information

## 1.1 S1 Appendix

Due to stochasticity, noise, and context-dependence (e.g. sequence-dependence) of the NAAB-amino acid interactions, a measurement performed on the *k*th target will yield an approximation 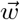 to the reference affinity vector 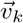. If we assume that the distribution according to which these measurements occur is Gaussian, then we can obtain a simple criterion for determining whether two N terminal amino acids will be distinguishable on the basis of affinity measurements made using a particular set of NAABs. We denote by 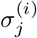 the standard deviation of the measurements made with NAAB *i* against amino acid *j*. For each amino acid, we may define a sphere of radius *ρ_j_*, centered on the vector 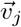, which surrounds that amino acid in affinity space. Here,

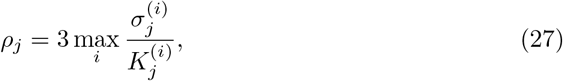

where 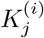 is the dissociation constant for the binding of the *i*th NAAB to the *j*th amino acid.

N-terminal amino acids will be identifiable with 99.9% certainty provided that there is no overlap in affinity-space between the *j* spheres of radius *ρ_j_*. To determine whether there is such an overlap, we must consider the distance metric

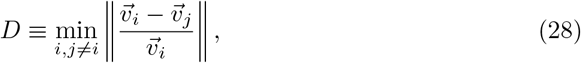

where the division is applied element-wise. In order to assign affinity measurements to the correct reference affinity 99.9% of the time, it is sufficient (but not necessary) to have

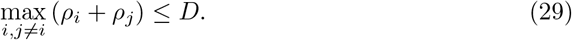

Using equation (27), it is then also sufficient to have

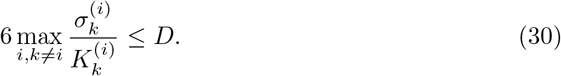

For the specific case of the NAAB affinity matrix, we find that *D* = 3.84. Thus, in order to ensure that the amino acids can be correctly identified 99.9% of the time, we must have

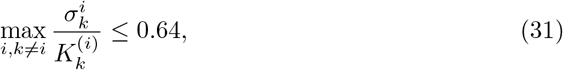

or, equivalently, the standard deviation of the *k_D_* measurements must be no greater than 64% of the mean.

## 1.2 S2 Appendix

Under the assumption of Poissonian noise, the photon rates in the bound and unbound states are given by

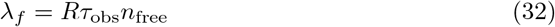

and

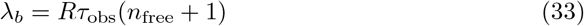

respectively. In order to be able to distinguish the bound state from the unbound state, it is clear that we must have

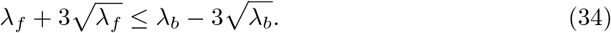

Because *λ_b_* > *λ_f_*, we may replace the standard deviation 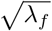 on the left-hand side by the standard deviation 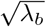, obtaining

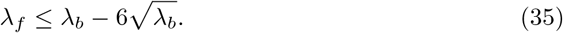

Hence,

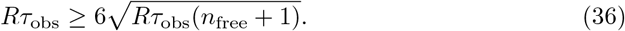

We find the final requirement:

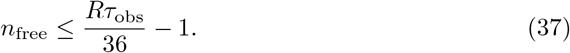

Rephrased as a condition on the concentration of the binder, we find

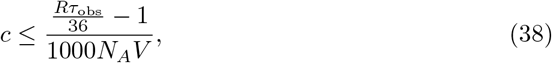

or

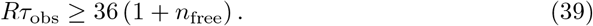

If *n*_free_ ≤ 1, then the assumption of Poissonian noise is invalidated because the emission of successive photons is not independent (it depends on the presence of fluorophores in the observation field). The assumption of Poissonian noise may also be invalidated if the frame rate is comparable to the rate at which fluorophores enter and leave the observation field. In either case, to correctly simulate the noise, one must draw the number of free binders that enter the observation field during a given frame from a Poisson distribution with mean *n*_free_*τ*_obs_/*τ*_dwell_, where *τ*_dwell_ is the amount of time each binder spends in the observation field on average. The average dwell time of free binders in a region of thickness Δ*x* may be calculated as

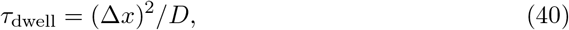

where *D* is the diffusion constant [23]. For a small protein in water, we have *D* ~ 10^−10^ m^2^ s^−1^. Taking Δ*x* = 100 nm, we find that free binders will dwell on average *τ*_dwell_ = 100 μs within the imaging plane.

Once the number of binders entering the observation field during the frame has been determined, one must draw the length of time *t* that each binder remains in the frame from an exponential distribution with mean *τ*_dwell_. Finally, for each binder, one must draw the number of photons emitted by that binder from a Poisson distribution with mean *Rt*. When the number of free binders is small, the resulting noise will differ significantly from Poisson noise due to the exponential distribution over dwell times. In our simulations, the long tail of the exponential distribution tends to significantly increase the difficulty of distinguishing transient binding and unbinding events, compared to simple Poisson noise (data not shown).

## 1.3 S3 Appendix

The intensity *I* is related to the photon rate *R* of the fluorophore by

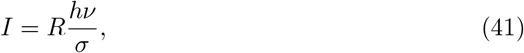

where *h* is Planck’s constant, *ν* is the frequency, *σ* is the absorption cross-section of the fluorescent dye, and *R* is the rate of absorption. To determine the cross-section, we note that from the Beer-Lambert law,

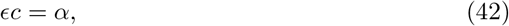

where *α* is the attenuation coefficient, *c* is the molar concentration, and e is the molar absorptivity, which we assume is given in M^−1^ m^−1^. Furthermore, we have

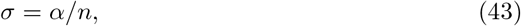

where *σ* is the absorption cross-section and *n* is the atomic number density. Hence, we have

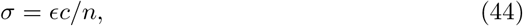

or, since *c* is the molar concentration and *n* is the number density, we have *n* = 1000*N*_*A*_*c*, where *N_A_* is Avogadro’s constant, *c* is given in molar and *n* is given in atoms per cubic meter. Thus,

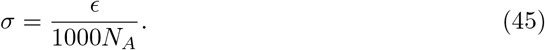

Hence, the photon number is given in terms of the intensity by

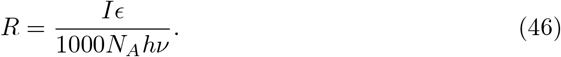

## 1.4 S4 Appendix

One advantage of occupancy measurements is that if *k*_on_ is known, then *k*_off_ may be determined even in the presence of photobleaching. To do so, we note that *T_i_* and *T_b_* are independent variables that depend on *k*_off_, *k*_on_, and *N_q_*. In the above analysis, we assumed that *N*_*q*_ was infinite, so that quenching could be neglected. If *N*_*q*_ is finite, however, then the true expressions for *T_i_* and *T_b_* are given by

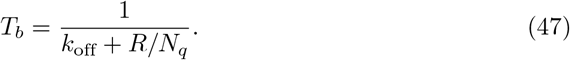

and

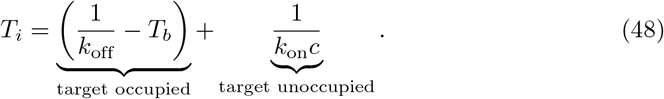

The first term in equation (48) is the average time the target spends occupied by a quenched fluorophore, while the second term is the average time the target spends unoccupied between unbinding and binding events. Hence, if *k*_on_ is known, then *k*_off_ and *N_q_* may be determined from *T_b_* and *T_i_*.

## 1.5 S5 Appendix

In contrast to occupancy measurements, luminosity measurements are sensitive to error in the calibration of the measurement apparatus. Calibration error arises from a combination of systematic differences in the brightness of the on- and off-states, which may result if different NAABs have different numbers of fluorophores on average, and from systematic error in the measurement of the brightnesses of the on- and off-states. Systematic variation in the brightnesses of the fluorophores can be overcome by calibrating the device prior to each measurement (as discussed below). In general, however, systematic error in the measurement of *S* and *N* significantly disrupts attempts to determine the absolute value of *k_D_* due to divergences in the derivative of *k_D_* as *M* approaches *N*. Hence, for weak binders in particular, infinitesimal changes in the calibration level can lead to divergent changes in the measured value of *k_D_*. For this reason, if the goal of the measurement is to determine the absolute value of *k_D_*, it is essential that the concentration be chosen such that the value of M to be measured lies close to *S*, i.e., such that the concentration *c* is close to or greater than *k_D_*. If *k_D_* is large or unknown, however, this requirement may not be achievable.

In our case, however, we are interested not in determining the absolute value of *k_D_*, but rather in determining the identity of a target (N-terminal amino acid) from the binding affinities of many binders (NAABs). In this case, one may significantly reduce the effects of calibration error by using the reference values of *k_D_* to calculate the expected photon rate *E* from the brightnesses of the on- and off-states, for each of the possible target identities. After having performed the measurement with all 17 binders, one is left with a vector 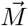 of the photon rates measured for each binder, and a set of vectors 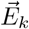, the *k*th of which is the vector of photon rates that one would have expected to measure if the target were of type *k*. The identity of the target is then determined by minimizing the norm of 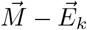 over *k*. The key difference here is that because one compares the expected photon rates to the measured photon rates, one avoids the nonlinearities inherent in calculating the measured dissociation constant from the measured photon rate.

**Fig. 6.**
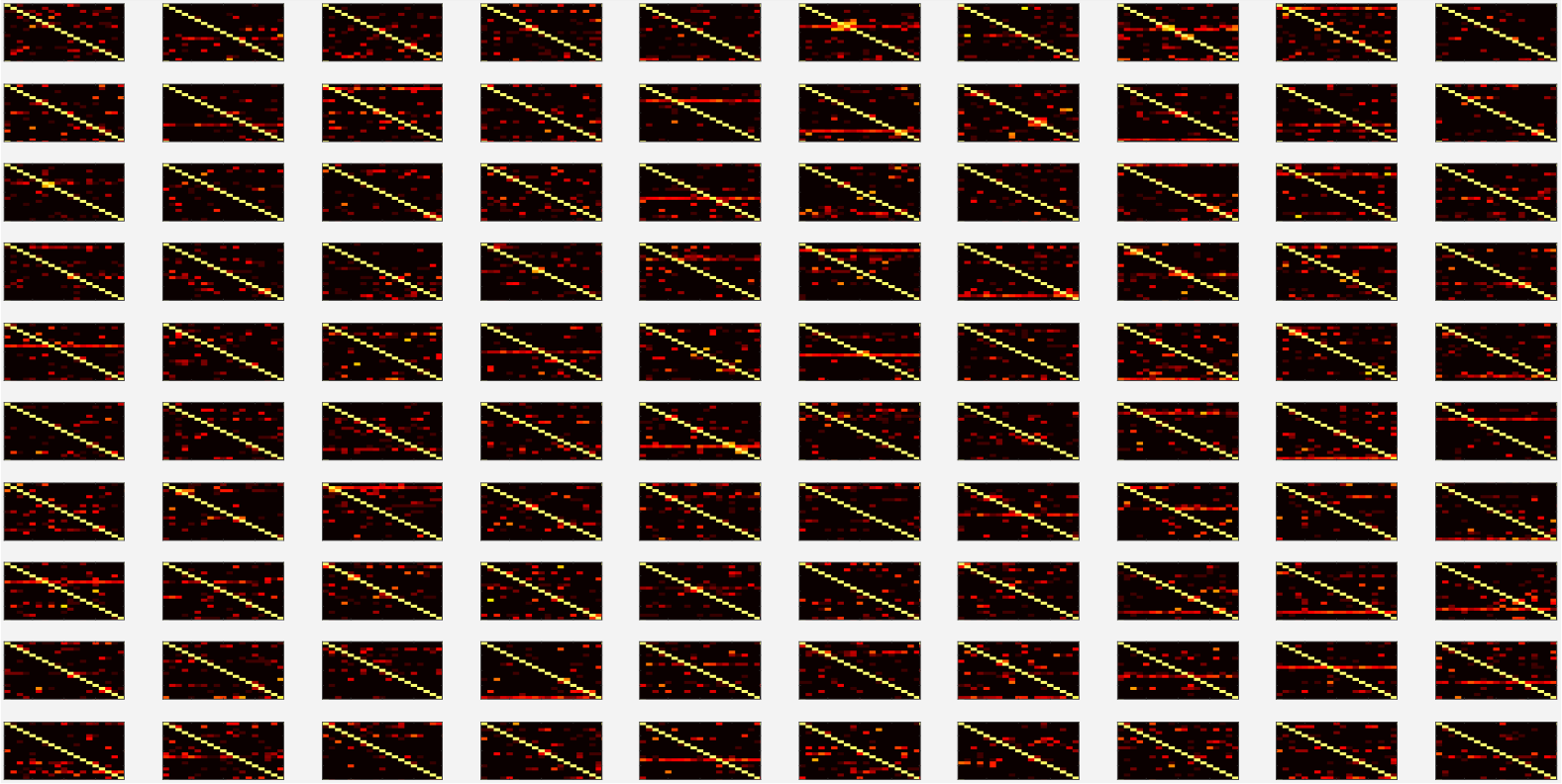
Accuracies for amino acid calling obtained for 100 random affinity matrices in simulations. 100 random affinity matrices were generated by randomly shuffling the entries of the NAAB affinity matrix. For each resulting matrix, we simulated 10000 amino acid calls, with 5% calibration error and 0.25% kinetic error. The resulting accuracy matrices are presented here. The scale and axes for each matrix are identical to those in fig. 4E.

## 1.6 S6 Appendix

Figure 6 shows the full set of accuracy matrices determined by simulation for 100 random affinity matrices.

## Acknowledgments

We thank Andrew Payne, Jim Havranek and Jeff Nivala for helpful discussions. SR acknowledges funding from the Fannie and John Hertz Fellowship. ESB acknowledges funding from NIH 1R01MH114031, John Doerr, Open Philanthropy, NIH 1R01EB024261, the HHMI-Simons Faculty Scholars Program, U.S. Army Research Laboratory and the U. S. Army Research Office under contract/grant number W911NF1510548, and NIH Director’s Pioneer Award 1DP1NS087724.

